## Supplementary Figures for "Pervasive Translation in Mycobacterium tuberculosis"

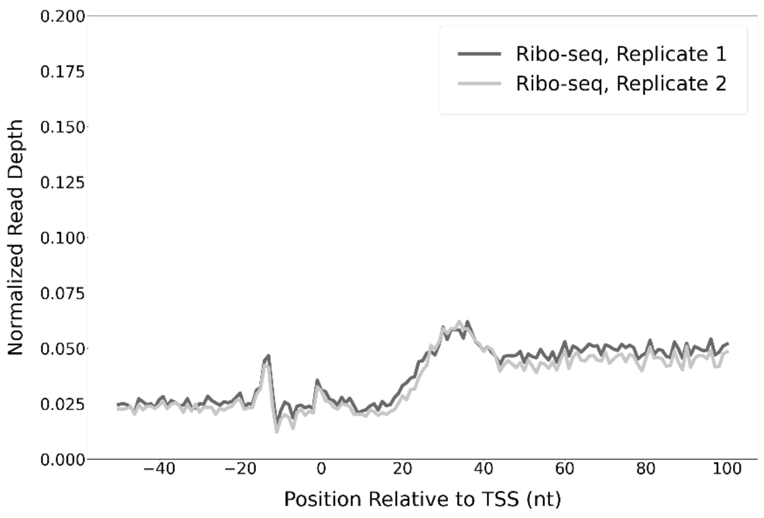


**Figure 1 - Figure Supplement 1. Modest enrichment of Ribo-seq signal downstream of the transcription start sites of non-leaderless RNAs.** Metagene plot showing normalized Ribo-seq sequence read coverage (data indicate the position of ribosome footprint 3' ends) in the region from -50 to +100 nt relative to the TSSs of RNAs that are not leaderless mRNAs.

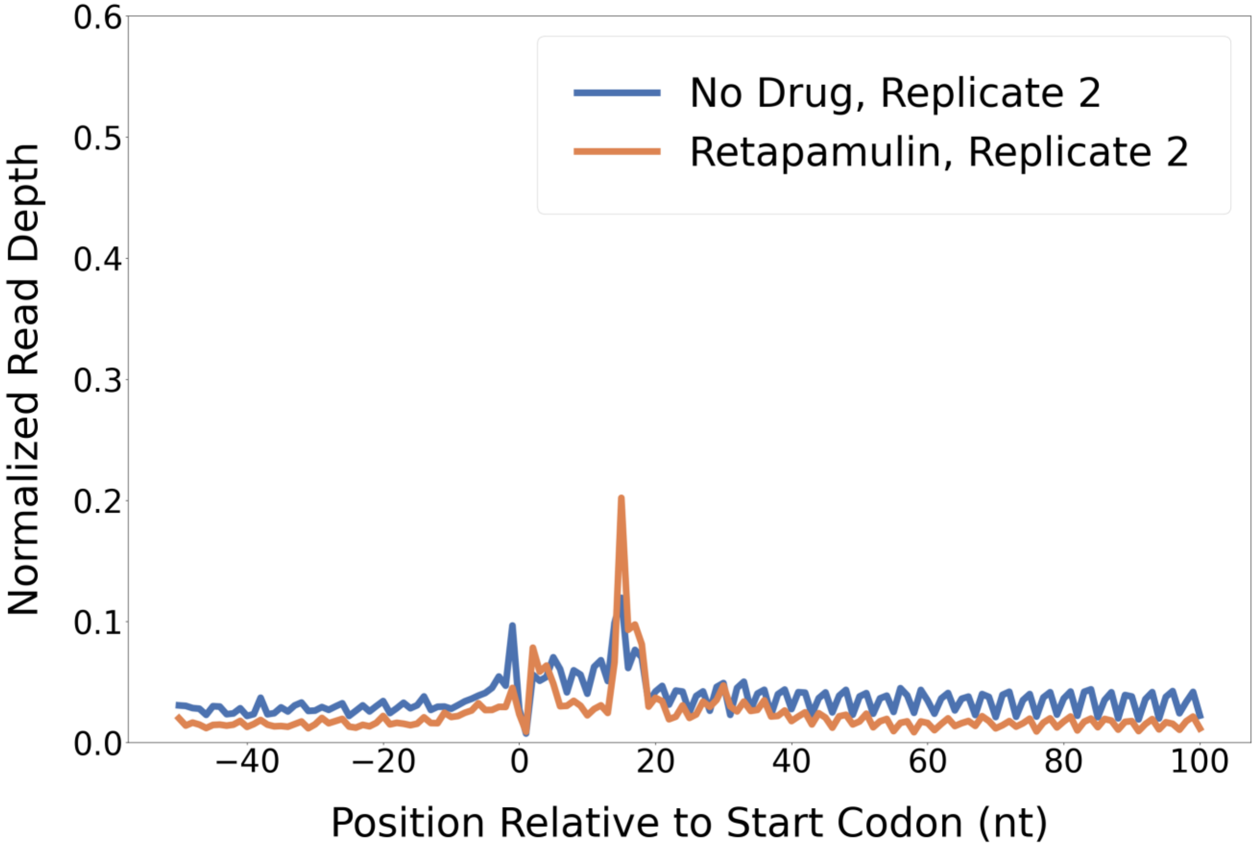


**Figure 2 - Figure Supplement 1. Retapamulin treatment traps initiating ribosomes.** Metagene plot showing normalized Ribo-seq and Ribo-RET sequence read coverage (single replicate for each; data indicate the position of ribosome footprint 3’ ends) in the region from -50 to +100 nt relative to the start codons of annotated, leadered ORFs. Figure 2A shows data for the other replicate datasets.

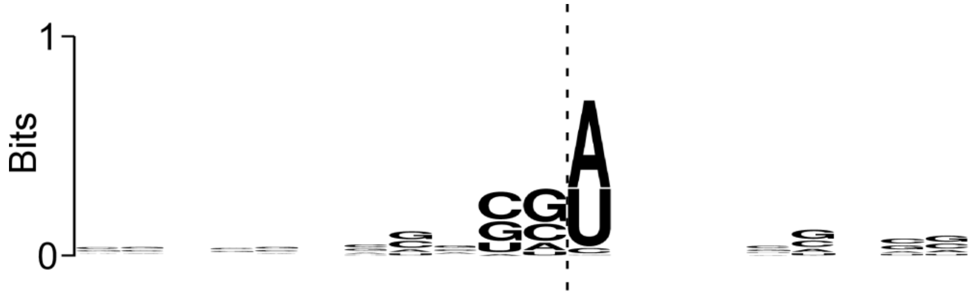


**Figure 2 - Figure Supplement 2. Sequence bias associated with the 3’ ends of ribosome-protected RNAs at IERFs.** Logo showing sequence bias around the 3’ ends of Ribo-RET RNA fragments associated with IERFs. The cleavage site at the 3’ end of the aligned RNA fragments is indicated by a vertical dashed line.

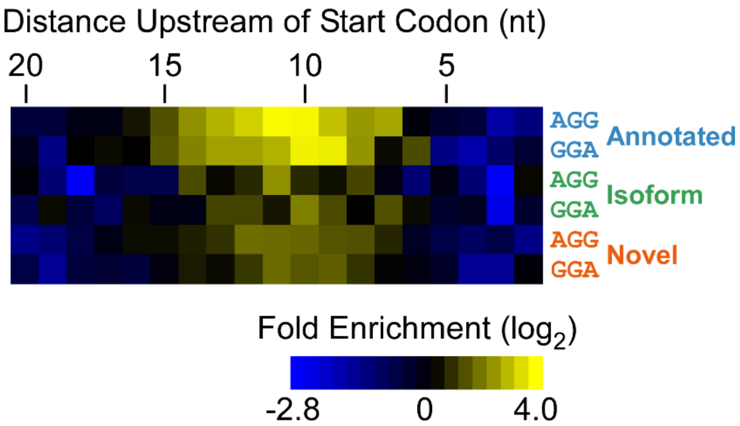


**Figure 3 - Figure Supplement 1. Enrichment of SD-like sequences upstream of higher-confidence ORFs identified by Ribo-RET.** Heatmap showing the enrichment of AGG and GGA trinucleotide sequences relative to control regions, for positions upstream of the start codons of annotated, isoform ORFs identified by Ribo-RET (higher-confidence set of ORFs)

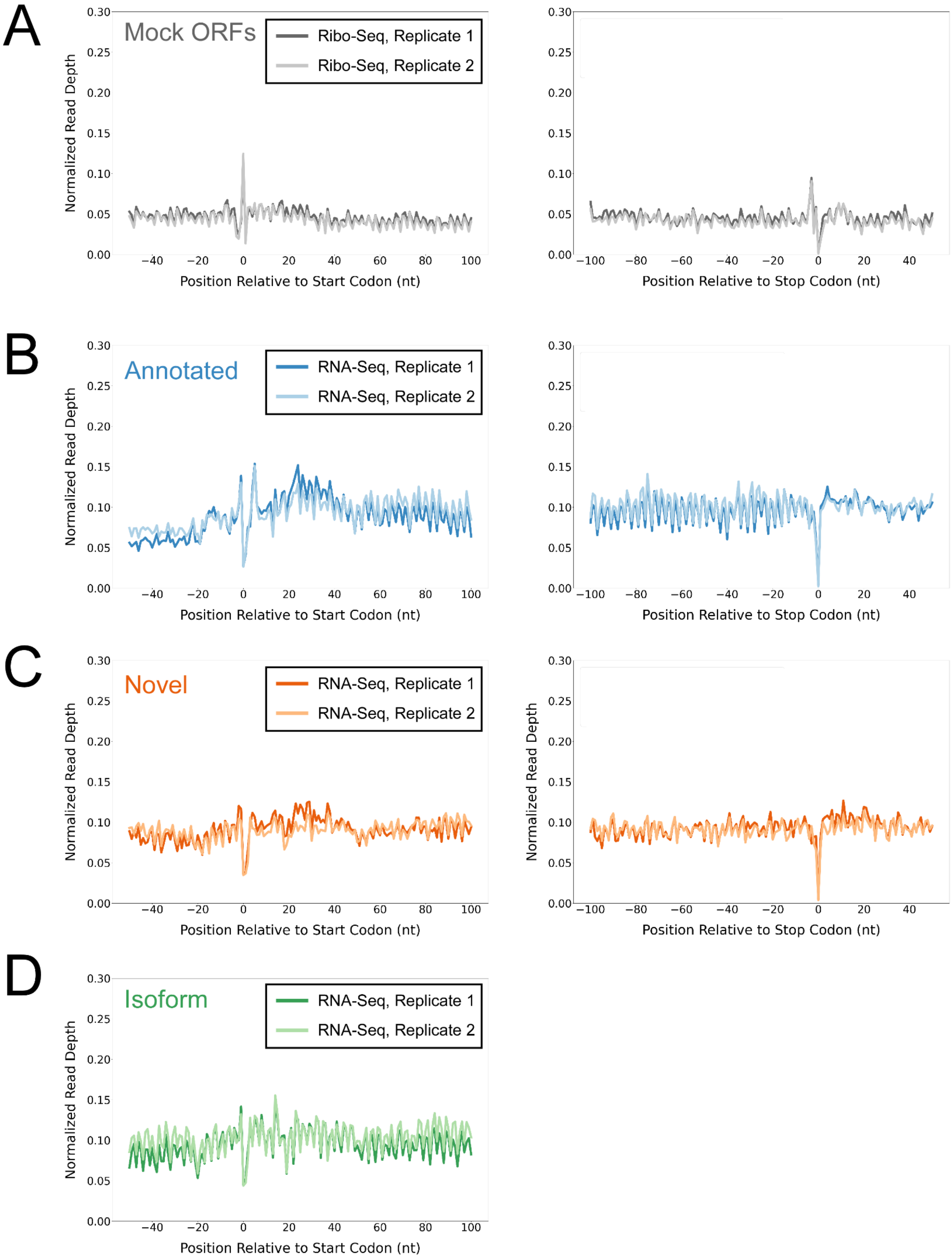

A
Mock ORFs
Ribo-Seq, Replicate 1
Ribo-Seq, Replicate 2
B
Annotated
RNA-Seq, Replicate 1
RNA-Seq, Replicate 2
C
Novel
RNA-Seq, Replicate 1
RNA-Seq, Replicate 2
D
Isoform
RNA-Seq, Replicate 1
RNA-Seq, Replicate 2
Normalized Read Depth
Position Relative to Start Codon (nt)
Position Relative to Stop Codon (nt)
0.00
0.05
0.10
0.15
0.20
0.25
0.30
−100
−80
−60
−40
−20
0
20
40
60
80
100

**Figure 4 - Figure Supplement 1. Control analyses using mock ORFs or RNA-seq data. (A)** Metagene plot showing normalized Ribo-seq sequence read coverage (data indicate the position of RNA fragment 3’ ends) for untreated cells in the regions around start (left graph) and stop codons (right graph) of mock ORFs. **(B)** Metagene plot showing normalized RNA-seq sequence read coverage (read 3’ ends) for untreated cells in the regions around start (left graph) and stop codons (right graph) of annotated ORFs identified from Ribo-RET data. **(C)** Equivalent data to **(B)** but for putative novel ORFs identified from Ribo-RET data. **(D)** Equivalent data to **(B)** but for putative isoform ORFs identified from Ribo-RET data. Only data for start codons are shown because the same stop codon is used by both an annotated and isoform ORF.

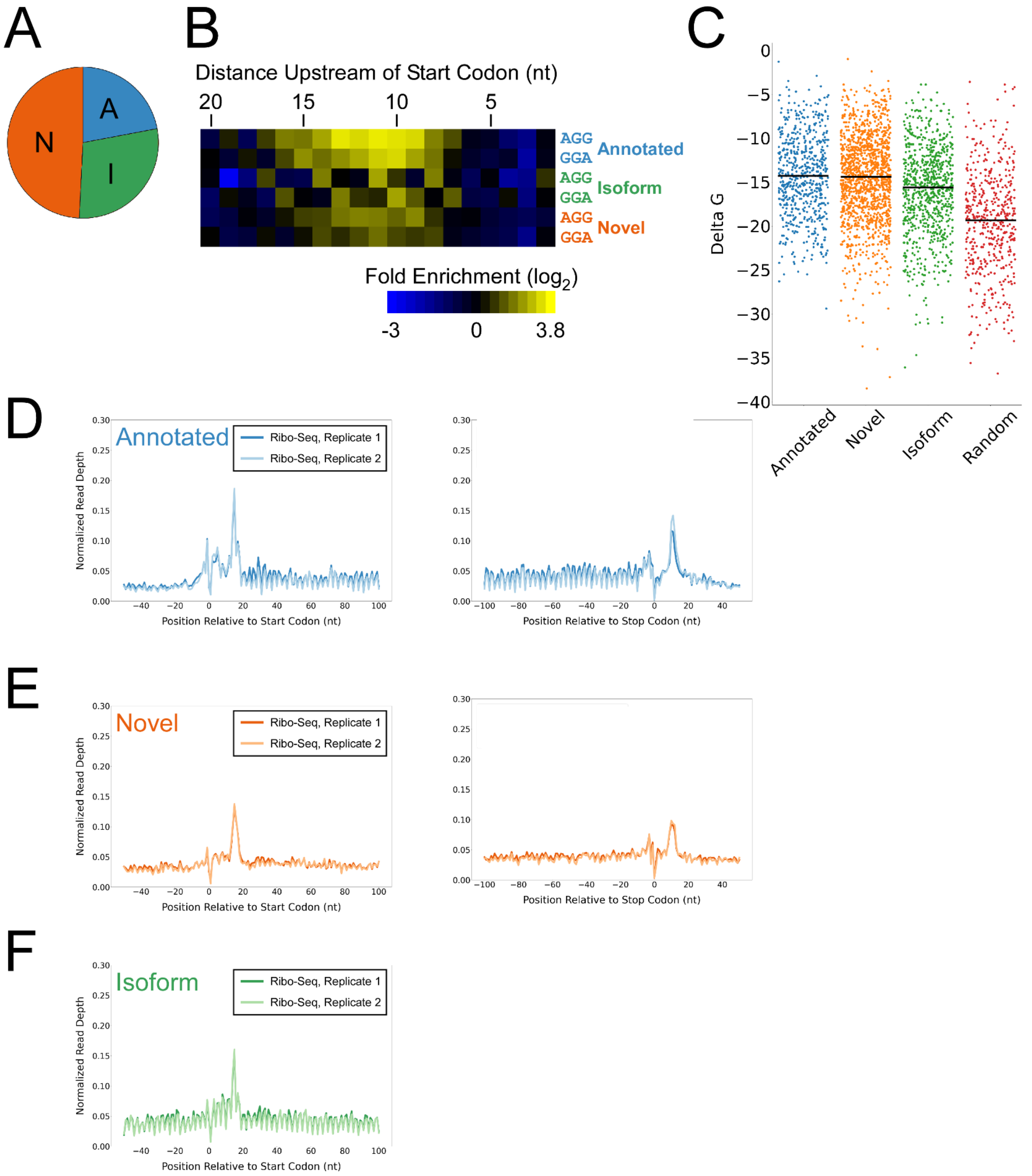
A
N
A
I
B
Distance Upstream of Start Codon (nt)
20
15
10
5
AGG
GGA
Annotated
AGG
GGA
Isoform
AGG
GGA
Novel
Fold Enrichment (log2)
-3
0
3.8
C
Delta G
Annotated
Novel
Isoform
Random
D
Annotated
Ribo-Seq, Replicate 1
Ribo-Seq, Replicate 2
Normalized Read Depth
Position Relative to Start Codon (nt)
Position Relative to Stop Codon (nt)
E
Novel
Ribo-Seq, Replicate 1
Ribo-Seq, Replicate 2
Normalized Read Depth
Position Relative to Start Codon (nt)
Position Relative to Stop Codon (nt)
F
Isoform
Ribo-Seq, Replicate 1
Ribo-Seq, Replicate 2
Normalized Read Depth
Position Relative to Start Codon (nt)

**Figure 4 - Figure Supplement 2**. **Features of lower-confidence ORFs identified by Ribo-RET. (A)** Distribution of different classes of lower-confidence ORFs identified by Ribo-RET. **(B)** Heatmap showing the enrichment of AGG and GGA trinucleotide sequences relative to control regions, for positions upstream of the start codons of lower-confidence annotated, isoform, and novel ORFs identified by Ribo-RET. **(C)** Strip plot showing the ΔG for the predicted minimum free energy structures for the regions from -40 to +20 nt relative to start codons for the different classes of lower-confidence ORF, and for a set of 500 random sequences. Median values are indicated by horizontal lines. **(D)** Metagene plot showing normalized Ribo-seq sequence read coverage (data indicate the position of ribosome footprint 3' ends) for untreated cells in the regions around start (left graph) and stop codons (right graph) of lower-confidence annotated ORFs identified from Ribo-RET data. **(E)** Equivalent data to **(D)** but for lower-confidence isoform ORFs identified from Ribo-RET data. Only data for start codons are shown because the same stop codon is used by both an annotated and isoform ORF. **(F)** Equivalent data to **(D)** but for lower-confidence novel ORFs identified from Ribo-RET data.

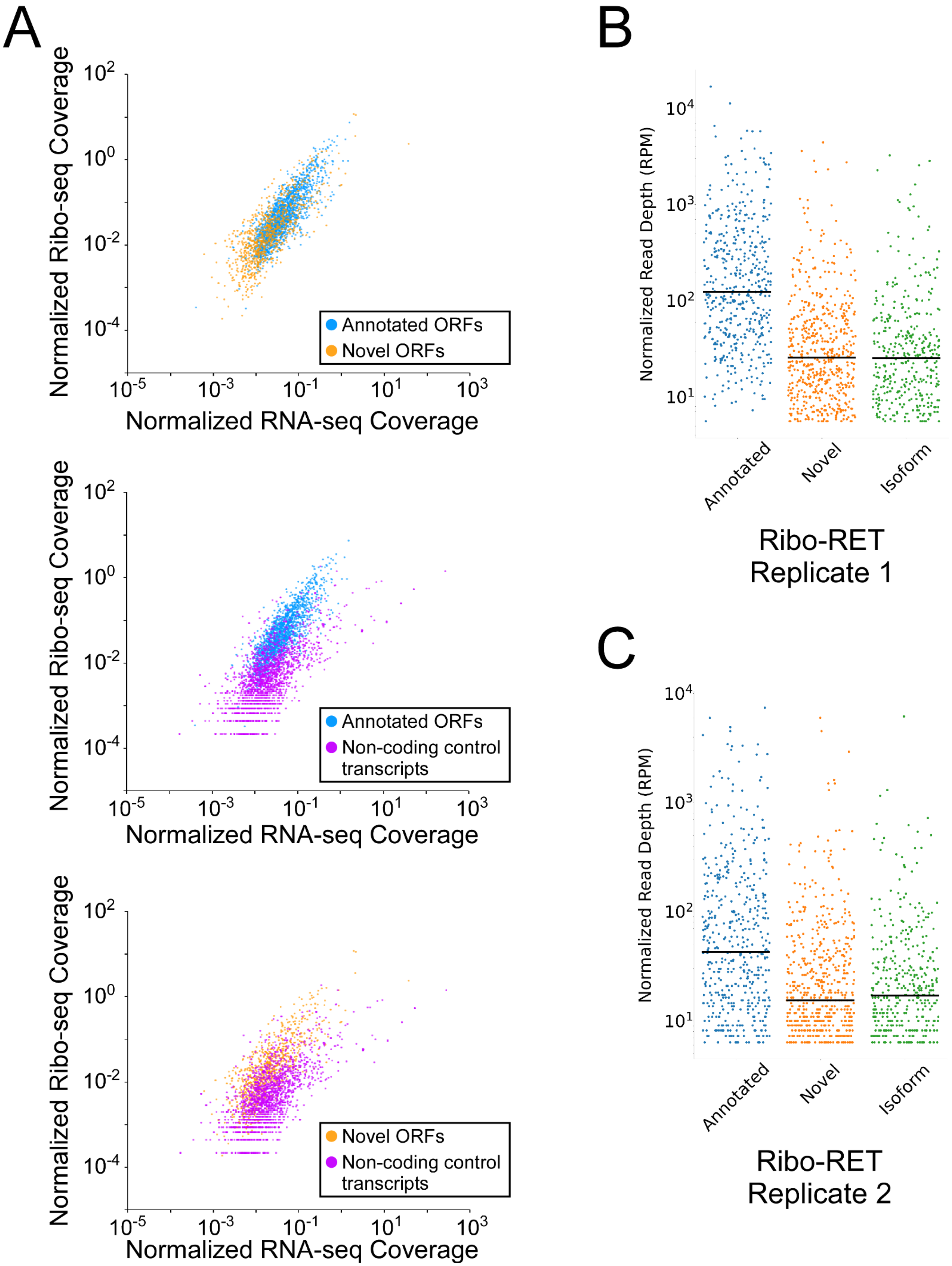
A
Normalized Ribo-seq Coverage
Normalized RNA-seq Coverage
Annotated ORFs
Novel ORFs
Non-coding control transcripts
B
Normalized Read Depth (RPM)
Annotated
Novel
Isoform
Ribo-RET Replicate 1
C
Ribo-RET Replicate 2

**Figure 5 - Figure Supplement 1. Novel and isoform ORFs are expressed at lower levels than annotated ORFs. (A)** Pairwise comparison of normalized RNA-seq and Ribo-seq coverage for annotated, novel and non-coding control transcripts. Reads are plotted as RPM per nucleotide using a single replicate of each dataset for reads aligned to the reference genome at their 3' ends (c.f. Figure 5, which shows data for the other replicate for each dataset). The categories compared are: (i) annotated ORFs (higher-confidence and lower-confidence ORFs detected by Ribo-RET, and leaderless ORFs; blue datapoints), (ii) novel ORFs (higher-confidence and lower-confidence ORFs detected by Ribo-RET and leaderless ORFs, for regions at least 30 nt from an annotated gene; orange datapoints), and (iii) a set of 1,854 control transcript regions that are expected to be non-coding (see Methods; purple datapoints). ORF/transcript sets are plotted in pairs to aid visualization. **(B)** Strip plot showing the normalized sequencing read depth for a single Ribo-RET replicate dataset, at start codons of higher-confidence annotated, isoform, and novel ORFs identified by Ribo-RET. Median values are indicated by horizontal lines. **(C)** Equivalent to **(B)** but for a second replicate Ribo-RET dataset.
